# Incorporation of macrophage immune stresses into an assay for drug tolerance in intracellular *Mycobacterium tuberculosis*

**DOI:** 10.1101/2025.05.09.653069

**Authors:** Greana Kirubakar, Luana Johnston, Bom Nae Rin Lee, David G. Russell, Nelson V. Simwela

## Abstract

Development of new and improved tuberculosis (TB) chemotherapies is hampered by antibiotic resistance and drug tolerance by *Mycobacterium tuberculosis* (*Mtb*). Phenotypic drug tolerance, a phenomenon where *Mtb* populations can temporarily survive therapeutic antibiotic concentrations, represents a significant hurdle to TB treatment and is indeed one of the factors responsible for prolonged TB therapy. Assays that can identify compounds with improved efficacy against drug tolerant *Mtb* are urgently required to improve TB treatment regimens. Here, we report the development of a 96-well plate assay capable of identifying anti-*Mtb* drugs with activity against drug tolerant *Mtb* in physiologically relevant intracellular environments within macrophages. Primary murine macrophages modified either by immunological activation or specific CRISPR/Cas9 gene knockouts to generate tolerance-inducing environments were infected with an *Mtb* strain constitutively expressing luciferase. Following drug exposure, differences in bacterial survival were measured by bacterial outgrowth after lysis of the host macrophages. By monitoring *Mtb* luciferase in infected macrophages before, during and after drug treatment, we confirmed earlier observations that host immune stresses trigger induction of drug tolerance. However, while host stresses induced tolerance against some anti-TB compounds, the same host stresses were synergistic with other anti-TB drugs. Our assay provides the ability to profile the activities of anti-TB drugs on bacteria in intracellular host environments which is critical to the rational design of drug combinations that provide optimal coverage of the *Mtb* sub-populations in the infected host.

**Author summary:** TB treatment is a lengthy process which at minimum takes 6-9 months in cases of drug sensitive *Mtb*. Long TB treatments are in part, due to drug tolerance phenotypes in the bacterial population which can be a result of immune related stresses on the bacteria in the infected host cells. We used this knowledge to develop a luciferase-based assay which can be used to screen, optimize and discriminate anti-TB drugs with enhanced activity against drug tolerant *Mtb* inside the infected host cells. Primary murine macrophages were modified either by cytokine activation or genetic knockout of certain host genes to mimic immune related stresses experienced by the bacteria *in vivo*. By infecting the modified macrophages with an *Mtb* strain expressing luciferase followed by drug addition, our assay was able to confirm that immune related stress induce drug tolerance to *Mtb*, and that the drug tolerance phenotypes are unique to certain drug classes consistent with their known mode of action. The assay provides an important addition to anti-TB drug discovery by providing a means to readily screen new drugs and drug combinations that display improved activity in tolerance inducing environments in a physiologically relevant context.

## Introduction

*Mycobacterium tuberculosis* (*Mtb*) remains the single most frequent cause of death by an infectious agent. The current treatment regimen for tuberculosis (TB) commonly comprises of four drugs, Isoniazid (INH), Rifampicin (RIF), Pyrazinamide (PZA) and Ethambutol (EMB) administered for a period of 6-9 months in the case of drug-sensitive *Mtb* [1]. The major driver behind this prolonged treatment is the differential drug susceptibility across the bacterial population *in vivo*.

Differential drug susceptibility in bacteria is usually the product of three distinct phenomena; persistence, phenotypic tolerance and mutation [2, 3]. The first two responses are non-heritable, while acquisition of drug resistance by genetic mutations spreads clonally across the bacterial population. Persistence in the face of drug pressure is usually exhibited by a very small number of bacteria capable of surviving lytic concentrations of antibiotic and will re-enter the growth phase upon removal of drug pressure. The number of these persistors remains constant upon rechallenge and the mechanism(s) behind this property remain elusive. Phenotypic drug tolerance, on the other hand, is distinct from persistence and can be shown by a large proportion of the bacterial population at any given time. *In vitro* studies on *Mtb* have revealed multiple mechanisms linked to environmental stress or the generation of heterogeneity within the bacterial population that can lead to phenotypic drug tolerance [4-11]. We believe that it is this phenomenon that is the major hurdle to shortening drug treatment regimens for tuberculosis, and one of the most significant factors in the frequent emergence of heritable drug resistance in this pathogenic bacterium.

Several studies have shown how the immune environment *in vivo* can be a powerful inducer of phenotypic drug tolerance [12-15]. In one such study, *Mtb*-infected macrophages were flow-sorted from dissociated infected mouse lung into activated (CD80 hi) and resting (CD80 lo) host cells, established in culture overnight and exposed *ex vivo* to INH or RIF. When surviving bacteria were quantified by colony forming units (CFUs), it was found that the bacteria in the activated host macrophages exhibited greater survival [14]. Similar data were also generated in a single cell RNA-seq study conducted on mice infected with fluorescent fitness reporter *Mtb* demonstrating that *Mtb* under immune stress from immune-activated host macrophages were less drug susceptible, to INH and RIF at least [16]. More recently, Lavin and Tan linked comparable *Mtb* fitness and immune activation data to additional physiological host cell readouts and overlaid these data onto the topographical organization of the mouse lung granuloma in mice treated with INH, RIF and PZA [17]. Their results indicated that those bacteria in cells abutting the necrotic core of the granuloma, which appeared to be more actively replicating, were also more drug susceptible than the bacilli in macrophages at the periphery of the granuloma, where the host cells appeared more immunologically activated. All of these data stress the importance of incorporating host immune pressures in designing effective anti-TB drug regimens, a component that is rarely included in current drug discovery pipelines.

While attempts have been made to mimic the changing environments within macrophages by modulation of *Mtb* growth in broth media with respect to ionic content, pH, carbon source and oxygen tension [5, 7, 9, 11], experiments in growth media lack the environmental complexities encountered by the pathogen within a host and yield a limited understanding of drug tolerant bacteria and their vulnerabilities during infection [2]. Moreover, even though studies on macrophage heterogeneity *in vivo* have demonstrated how host cell physiology is a major driver of bacterial fitness, growth and drug susceptibility [18], the incorporation of host cell physiology into drug discovery or drug regimen design has been minimal. Using primary murine macrophages derived from conditionally immortalized myeloid percussors transduced with the estradiol-responsive Hoxb8 transcription factor [19], we have developed a 96 well plate-based assay that facilitates assessment of anti-TB drug efficacy in host macrophages modified either by immunological activation, or CRISPR/Cas9 gene editing to realign host cell metabolism and physiology. The results confirmed earlier conclusions that host immune stresses do trigger induction of drug tolerance, but that this tolerance differs with the identity or mode-of-action of the anti-TB drugs. We believe such data to be a critical component in the design of combinatorial drug regimens that provide optimal coverage of the heterogeneous bacterial sub-populations present in the infected host.

## Results and discussion

### Assay design and outline

Our assay involves the use of primary murine bone marrow derived macrophages (BMDM^h^s) derived from Hoxb8 conditionally immortalized myeloid progenitors [19]. BMDM^h^s retain similar characteristics to mouse primary bone marrow macrophages and are readily infected with *Mtb* [20-22]. Resting or activated (LPS/IFNγ) BMDM^h^s are infected with luciferase expressing *Mtb* (*Mtb* Erdman-Lux) at a multiplicity of infection (MOI) of 0.4. Anti-TB drugs are added 4 days post infection to allow for sufficient impact of immune stresses on the bacteria. In addition, BMDM^h^s in which CRISPR-Cas9 had been used to edit host genes resulting in *Mtb* growth restriction can also be used to induce stress in the bacteria. Luciferase measurements are made on day 0 and day 4 before drug addition and on days 5-8 post drug addition. After the day 8 luciferase read, macrophage media containing drugs is removed and the cell monolayers are gently washed with fresh media without drug at least once. *Mtb* growth media (7H9 broth) is then added to the wells to facilitate macrophage lysis. Bacterial recovery and outgrowth post-drug treatment is then measured by further luciferase reads on days 10 and 12 (assay outline summarized in Fig. 1).

**Fig. 1:**
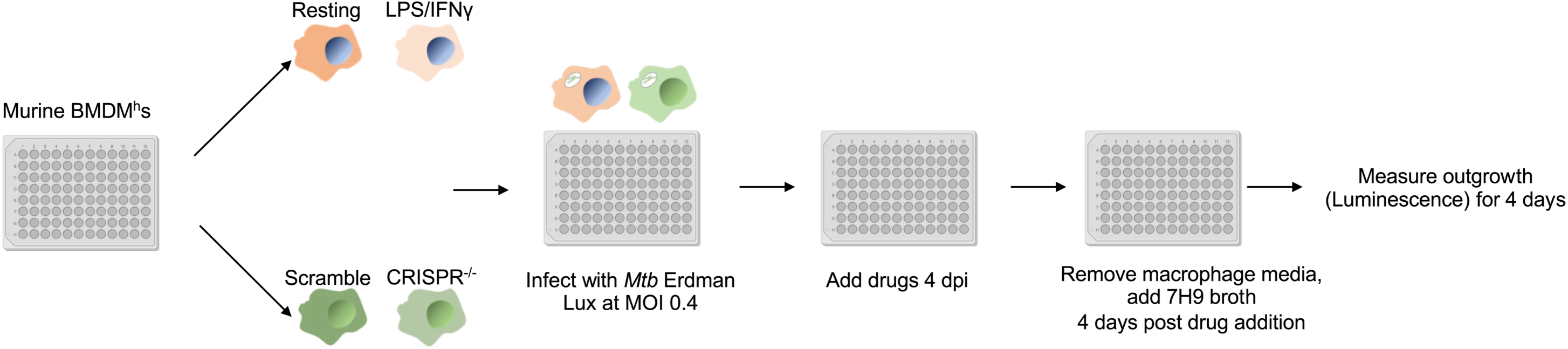
Assay workflow.

### LPS/IFNγ activation of BMDM^h^s induces drug tolerance to INH, RIF and EMB but not PZA

We had previously shown that increasing host antimicrobial immune pressure in BMDMs by LPS/IFNγ activation induces *Mtb* antibiotic tolerance to INH, RIF, EMB and PZA [14]. This quantitation was based on bacterial CFUs 48- or 96-hours post-infection in *Mtb* infected resting or activated BMDMs where drug treatment was commenced 2-hours post-infection. In the current assay, we infected resting or LPS/IFNγ activated wild-type BMDM^h^s (C57B6/J) with the *Mtb* Erdman-Lux strain at MOI 0.4 and allowed the infection to establish for 4-days. Extended infection prior to drug treatment enables the infection to establish and the host cell to robustly generate an intracellular environment that is nitrosative, oxidative and nutrient limiting in activated macrophages [23-25] in a manner we believe is representative of *in vivo* conditions. 96-well plate-based luciferase readouts increased the assay throughput and robustness as the measurements are non-destructive and bacterial growth kinetics can be monitored before, during and after drug treatment. Incorporation of a bacterial outgrowth readout through the addition of 7H9 broth to induce macrophage lysis further enabled the assay’s ability to identify drugs with improved activity against *Mtb* residing in drug tolerance promoting environments by measuring bacterial recovery when drug pressure is removed. In the absence of drugs, LPS/IFNγ activation significantly impacted *Mtb* growth in BMDM^h^s when compared to resting macrophages when we measured luciferase relative light units (RLU) on day 4 (Fig. 2). This is consistent with literature that immune activation is necessary for macrophages to maximize their tuberculostatic or tuberculocidal properties [25-27]. Activated BMDM^h^s induced observable stresses on the bacteria as early as 3 hours post-infection on day 0 (Fig. 2). Subsequently, four first-line anti-TB drugs INH, RIF, EMB and PZA were added on day 4 and bacterial growth was monitored up to day 8 when macrophage media was replaced with 7H9 broth. In comparison with resting macrophages, bacteria in activated BMDM^h^s displayed a reduced susceptibility to INH, RIF and EMB as evidenced by robust bacterial outgrowth on days 10 and 12 (Fig 2A-C). The degree of phenotypic drug tolerance to all the 3 drugs was evident at all the concentrations tested, however, lower concentrations resulted in significantly higher differential drug susceptibility. In contrast, *Mtb* in activated BMDM^h^s was more susceptible to PZA at 200 μg/ml or 800 μg/ml when compared to resting macrophages (Fig. 2D). INH and EMB act by inhibiting the synthesis of the *Mtb* cell wall. INH is a prodrug that is converted by the catalase-peroxidase KatG into an INH-NAD adduct which binds the NADH-dependent fatty acid synthase II (FAS-II) component InhA to inhibit mycolic acid synthesis [28-30]. EMB is a bacteriostatic agent which disrupts the production of the mycobacterial cell wall constituent, arabinogalactan [31] and may act in synergy with INH to inhibit InhA [32]. RIF exerts its bactericidal action by inhibiting the bacterial RNA polymerase B subunit to prevent RNA transcription. In activated macrophages, intracellular bacilli upregulates genes required for survival in stressed intracellular environments like acidic pH, nutrient limitation, hypoxia and exposure to reactive oxygen and nitrogen species. Concomitantly, expression of genes required for bacterial growth such as ribosomal biogenesis, amino acid biosynthesis, lipid breakdown and oxidative respiration are usually downregulated. This adaptation process remodels *Mtb* physiology to attain a metabolically quiescent state which renders the bacterium less susceptible to drugs like INH, EMB and RIF that are usually regarded as most active against replicating bacteria [13, 33]. Our outgrowth component of the assay is able to clearly resolve these subtle differences in drug susceptibility as upon drug removal and macrophage lysis in 7H9 broth on day 8, we observed a rapid amplification of luciferase signal from bacteria in activated macrophages incubated with INH, EMB and RIF when compared to *Mtb* from drug treated resting macrophages (Fig 2A-C).

**Fig. 2:**
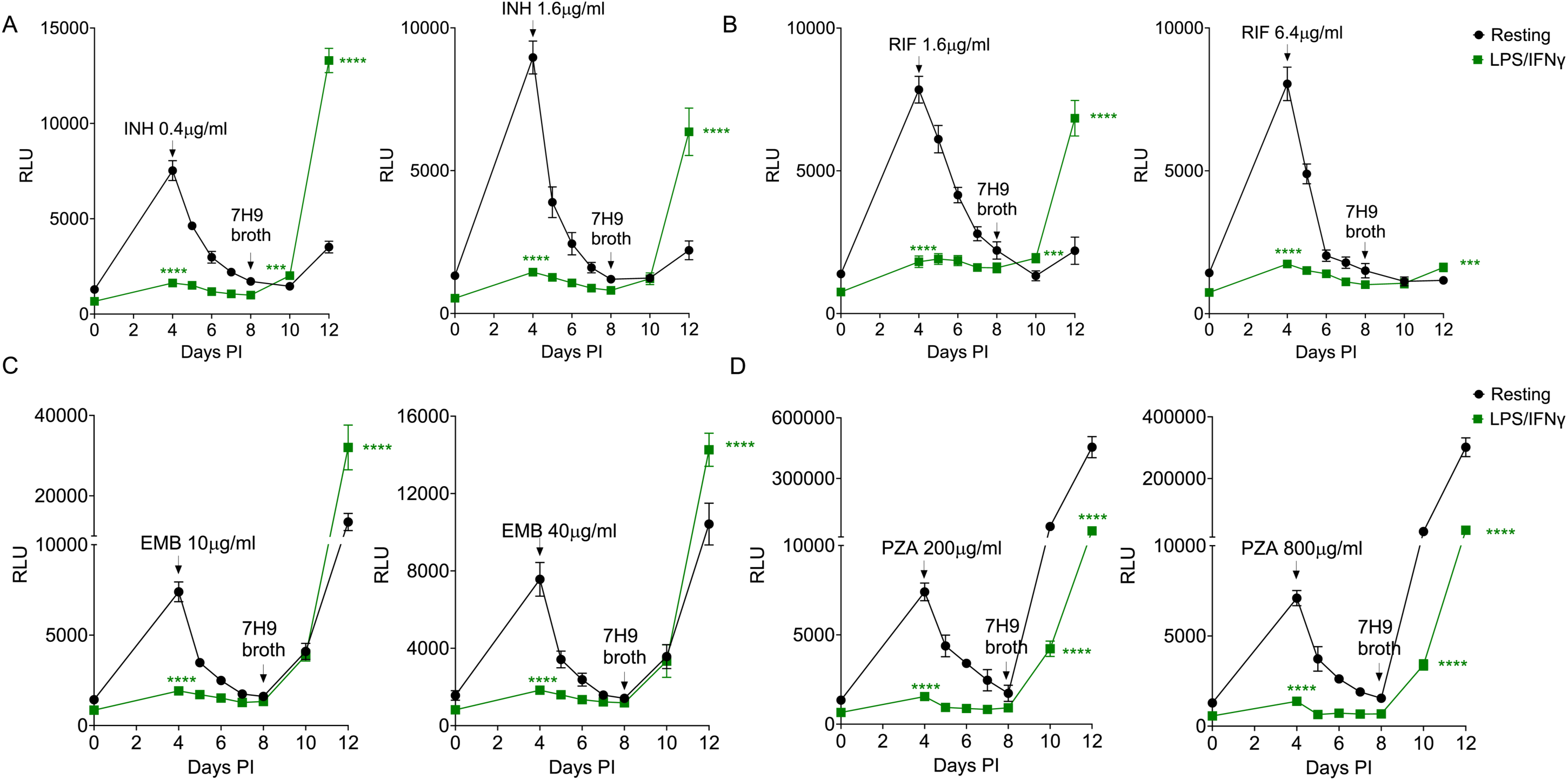
Mtb drug susceptibility to INH, RIF, EMB and PZA in LPS/IFNγ activated BMDM^h^s. Resting and activated macrophages were infected with the *Mtb* Erdman-Lux strain at MOI 0.4 for 4 days before addition of drugs. Drugs were added on day 4 (indicated by arrow). Macrophage media was replaced with 7H9 broth on day 8 (4 days after addition of the drugs). Bacterial growth was monitored by measuring luciferase relative light units (RLU). Growth kinetics of *Mtb* in macrophages treated with the indicated concentrations of INH (A), RIF (B), EMB (C) and PZA (D). n = 3 biological replicates. ****P < 0.0001, two-way ANOVA alongside Dunnett’s multiple comparison test. Data are presented as mean values ± SD.

PZA is a bacteriostatic prodrug that is converted to its active form pyrazinoic acid by a bacterial amidase pncA and results in widespread inhibition of cellular enzymes and energy production processes [34]. While our previous work suggested that *Mtb* acquires drug tolerance to PZA in activated BMDMs [14], our current data indicates that bacteria in activated BMDM^h^s were actually more susceptible to PZA than those in resting macrophages (Fig. 2D). We believe that prolonging of the duration of macrophage stress on the bacteria prior to drug exposure and the incorporation of the bacterial outgrowth readout significantly improves the resolution of our current assay. Given that PZA is most active in acidic environments [34, 35], macrophage activation, which enhances phagosome maturation and acidification [36, 37], would potentiate the activity of PZA and enhance its efficacy against *Mtb* in immune activated macrophage environments. More recent work by Lavin and Tan demonstrated that PZA, unlike INH or RIF, had broad activity against *Mtb* in necrotic granuloma lesions irrespective of the bacterial replication status [17]. This is consistent with our data that PZA appears more active against both replicating and non-replating bacteria and is enhanced in potency under macrophage induced stress conditions.

### Second line anti-*Mtb* drugs show enhanced activity against *Mtb* in LPS/IFNγ activated BMDM^h^s

We also used our assay to assess the activity of second line anti-TB drugs against drug tolerant bacteria in activated BMDM^h^s. We choose four compounds from the BPaLM combination drug regimen which are used to treat drug resistant TB [1]; Bedaquiline (BDQ), Pretomanid (Pa), Linezolid (LZD) and Moxifloxacin (MOX). BDQ, LZD and Pa displayed increased and better efficacy against *Mtb* in activated BMDM^h^s when compared to resting macrophages (Fig. 3A-C). The fluoroquinolone MOX showed increased activity against *Mtb* in both resting and activated macrophages as reflected by almost absent bacterial recoveries on days 10 and 12 (Fig. 3D). BDQ blocks ATP production by targeting the mycobacterial ATP synthase [38]. Pa is a narrow spectrum nitroimidazole which inhibits the production of *Mtb* mycolic acid and also acts directly as a nitric oxide (NO) donor to induce NO mediated respiratory poisoning within the bacterial cell [39, 40]. BDQ and Pa are reported to be active against both replicating and non-replicating bacteria [40, 41]. Our data are consistent with these reports as both BDQ and Pa had the greatest efficacy against *Mtb* in activated BMDM^h^s (Fig. 3A, 3C) amongst the four tested second-line TB drugs. LZD is a member of the oxazolidinone class of protein synthesis inhibitors and acts by blocking the initiation of protein synthesis [42]. Even though LZD is largely bacteriostatic and has the highest activity against *Mtb* in log phase, it has been reported to be active against metabolically quiescent, non-replicating *Mtb* [43]. This is consistent with our observations as LZD treatment resulted in robust bacterial outgrowth from both resting and activated BMDM^h^s with, however, bacteria in activated macrophages more significantly impacted (Fig. 3C). MOX is a broad-spectrum bactericidal antibiotic that functions by inhibiting DNA gyrase (topoisomerase II) and topoisomerase IV with marked potency against replicating *Mtb* [44-46]. MOX has also been shown to have activity against *Mtb* in a hollow-fiber environmental stress model [47]. In addition to inhibiting DNA gyrases which are part of housekeeping genes in most bacteria, MOX-mediated killing of *Mtb* also involves a respiratory downshift, redox-dependent stress and accumulation of reactive oxygen species [48]. Our data are consistent with the proposed broad-spectrum nature of MOX activity against *Mtb* as bacterial outgrowth was almost absent in both resting and activated BMDM^h^s (Fig. 3D).

**Fig. 3:**
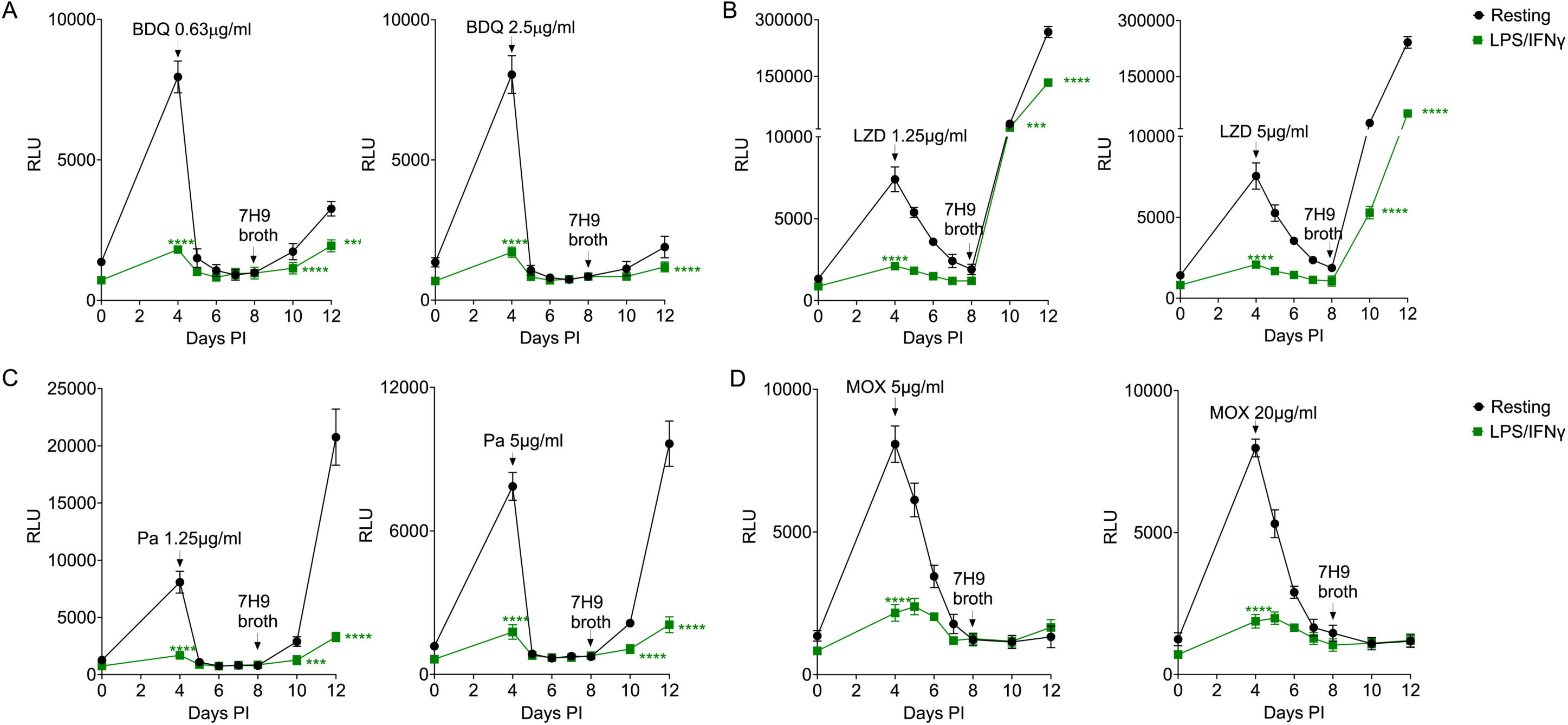
M*t*b drug susceptibility to BDQ, LZD, Pa and MOX in LPS/IFNγ activated BMDM^h^s. Macrophages were infected with bacteria and treated with drugs as in Fig.2. Growth kinetics of *Mtb* in BMDM^h^s treated with the indicated concentrations of BDQ (A), LZD (B), Pa (C) and MOX (D). n = 3 biological replicates. ***P < 0.001, ****P < 0.0001, two-way ANOVA alongside Dunnett’s multiple comparison test. Data are presented as mean values ± SD.

### GID/CTLH knockout BMDM^h^s elicit similar drug tolerance profiles to anti-TB drugs as LPS/IFNγ activated macrophages

In a recent CRISPR/Cas9 knockout screen, we found that macrophages lacking the mammalian GID/CTLH complex exhibited an enhanced ability to control the growth of intracellular bacteria [21]. The deletion of genes encoding members of the GID/CTLH complex activated a broad range of anti-microbial responses that resulted in oxidatively stressed and nutritionally starved *Mtb*. In LPS/IFNγ activated BMDM^h^s, the main inducer of the *Mtb* drug tolerance phenotype is NO mediated stress on the bacteria [14]. While NO is a strong anti-bacterial effector in murine macrophages, it has pleotropic impact on macrophage physiology and its route of induction in human macrophages is complex [49-54]. We sought to use GID/CTLH knockout macrophages as alternatives to LPS/IFNγ activated BMDM^h^s as they induce significant oxidative and nutritional stress on the bacteria but are strongly anti-inflammatory. As previously reported [21], infection of GID/CTLH knockout macrophages (*GID8*^-/-^, *MAEA*^-/-^) with *Mtb* Erdman-Lux resulted in significant reduction of RLUs by day 4 when compared to BMDM^h^s transduced with scrambled non-targeting control sgRNA (Fig. 4). In contrast to LPS/IFNγ activated BMDM^h^s, GID/CTLH knockout macrophages were less hostile to intracellular *Mtb* and supported some degree of bacterial expansion. However, we observed the induction of drug tolerance to the three first-line anti-*Mtb* drugs (INH, RIF, EMB) in GID/CTLH knockout macrophages (Fig. 4A-C) with outgrowth profiles comparable to those observed in LPS/IFNγ activated BMDM^h^s. Similarly, PZA was also more active against *Mtb* in GID/CTLH knockout macrophages when compared to control sgRNA (Fig. 4D). These data indicate that GID/CTLH knockout macrophages may offer an alternative host cell environment that leads to the induction of phenotypic drug tolerance in *Mtb* via NO-independent mechanisms. However, in contrast to LPS/IFNγ activated BMDM^h^s, we observed no differences in *Mtb* susceptibility to BDQ or Pa between GID/CTLH knockout macrophages and control sgRNA as the drugs were equally active against *Mtb* in all the conditions (Fig. 5A-B). Similarly, *Mtb* in GID/CTLH knockout macrophages showed a slight reduction in susceptibility to LZD and MOX (Fig. 5C-D). We believe this is a reflection of the differences in intensity of the experienced stresses by *Mtb* in LPS/IFNγ activated macrophages versus GID/CTLH knockout macrophages. *Mtb* in GID/CTLH macrophages exhibits reduced growth but is still replicating while activated macrophages almost completely block bacterial replication. Given that the magnitude of the bacterial outgrowth signal is also a function of bacteria population numbers at the time of host cell lysis, the outgrowth readouts in drug treated GID/CTLH knockout macrophages could therefore be slightly inflated when compared to scramble or LPS/IFNγ activated BMDM^h^s.

**Fig. 4:**
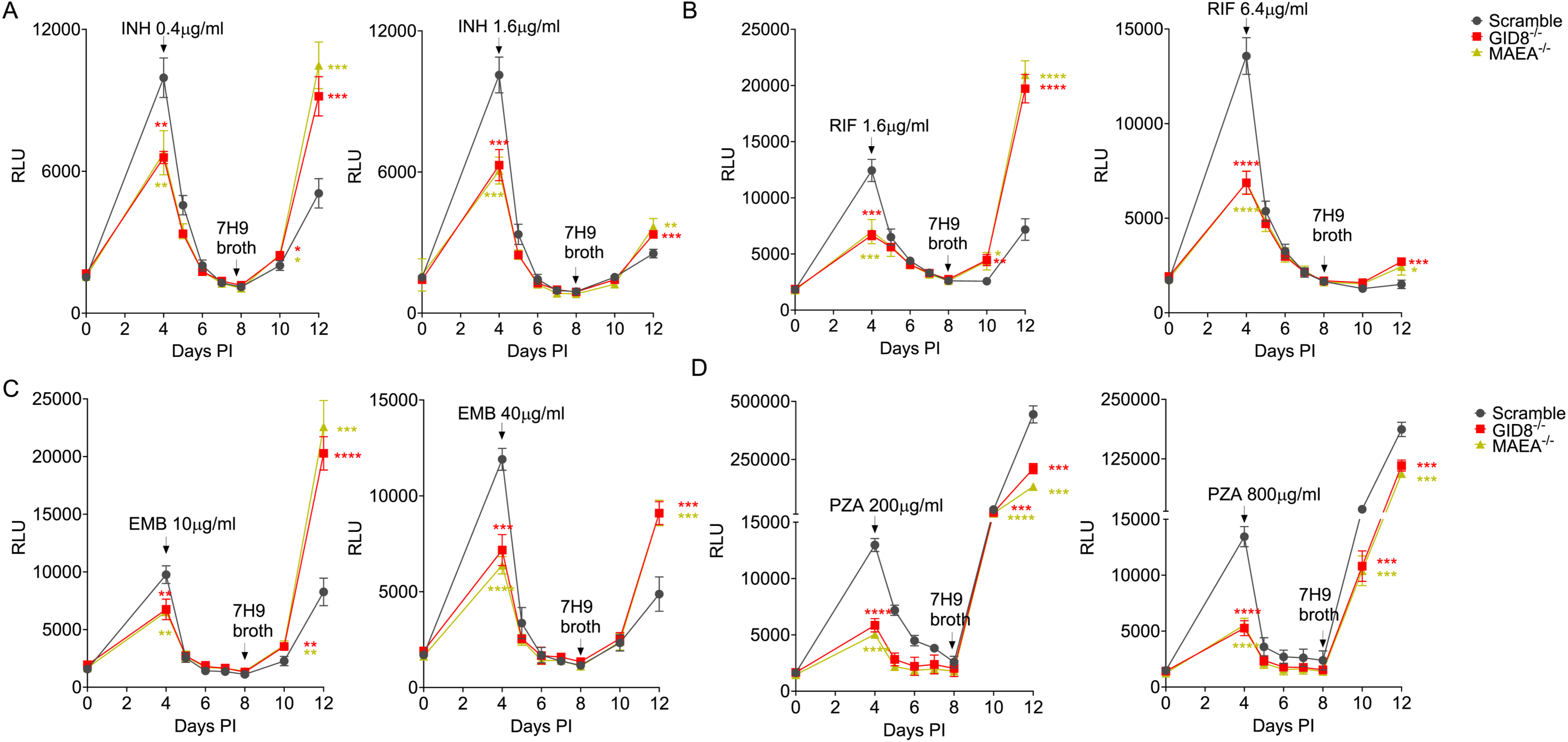
M*t*b drug susceptibility to INH, RIF, EMB and PZA in GID/CTLH knockout BMDM^h^s. Macrophages were infected with bacteria and treated with drugs as in Fig.2. Growth kinetics of *Mtb* in scramble or GID/CTLH knockout BMDM^h^s treated with the indicated concentrations of INH (A), RIF (B), EMB (C) and PZA (D). n = 4 biological replicates. *P < 0.01, **P < 0.01, ***P < 0.001, ****P < 0.0001, two-way ANOVA alongside Dunnett’s multiple comparison test. Data are presented as mean values ± SD.

**Fig. 5:**
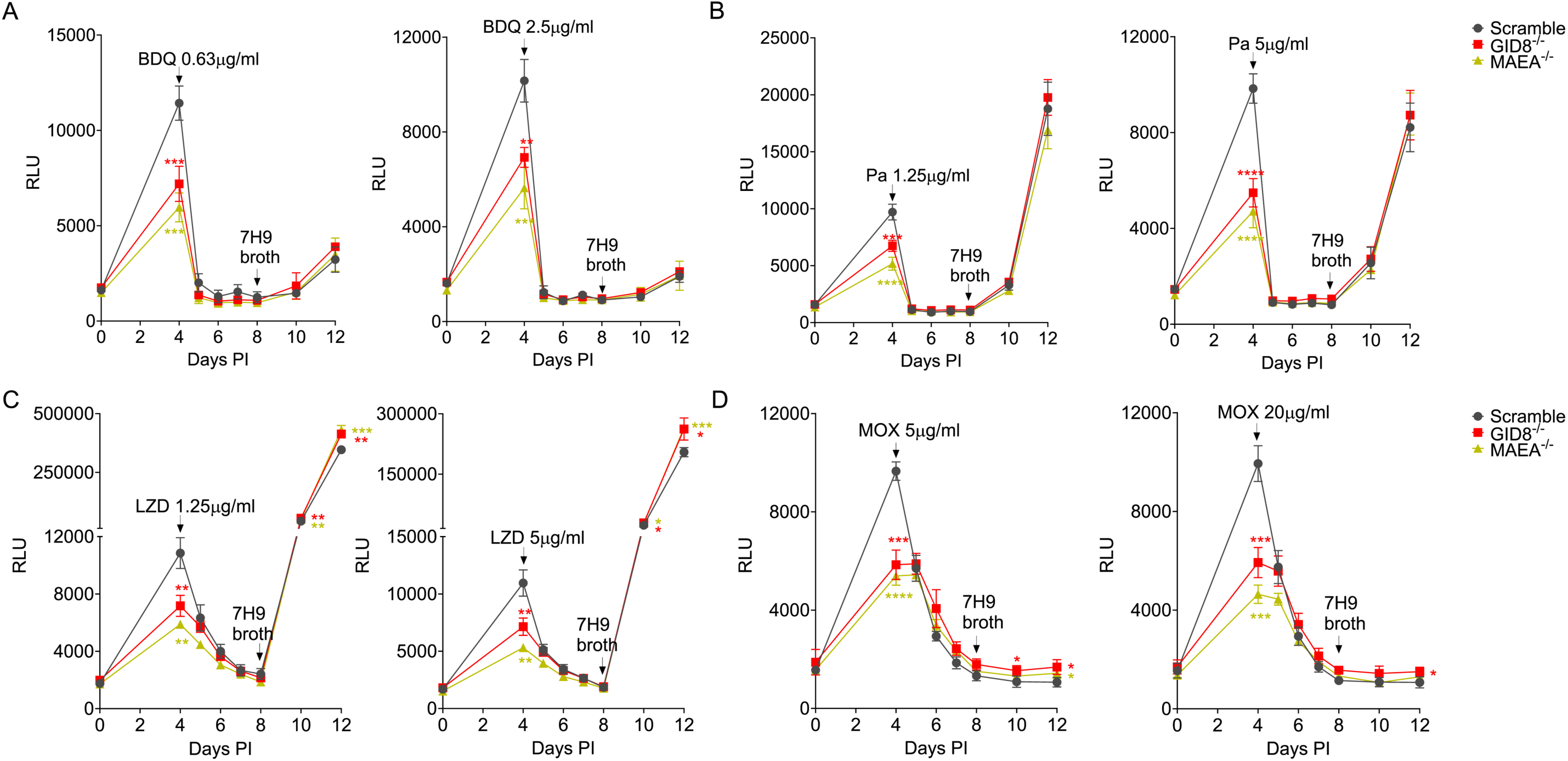
M*t*b drug susceptibility to BDQ, LZD, Pa and MOX in GID/CTLH knockout BMDM^h^s. Macrophages were infected with bacteria and treated with drugs as in Fig.2. Growth kinetics of *Mtb* in in scramble or GID/CTLH knockout BMDM^h^s treated with the indicated concentrations of BDQ (A), Pa (B), LZD (C) and MOX (D). n = 4 biological replicates. *P < 0.01, **P < 0.01, ***P < 0.001, ****P < 0.0001, two-way ANOVA alongside Dunnett’s multiple comparison test. Data are presented as mean values ± SD.

### Colony forming units (CFUs) measurements correlate with luciferase readouts in their ability to distinguish drugs with contrasting profiles in intracellular tolerance inducing environments

We next assessed whether drug tolerance profiles by luciferase readouts in our assay correlate with standard enumeration of bacterial CFUs upon drug treatment, which we had used previously [14]. To this end, we choose two drugs (INH and Pa) which displayed contrasting activity profiles in LPS/IFNγ activated and GID/CTLH knockout BMDM^h^s. Specifically, *Mtb* is highly tolerant to INH at 0.4 μg/ml in LPS/IFNγ activated and GID/CTLH knockout BMDM^h^s (Fig. 2A, Fig. 4A). To the contrary, Pa at 1.3 μg/ml is more effective against *Mtb* at least in LPS/IFNγ activated BMDM^h^s (Fig. 3C). We infected resting, LPS/IFNγ activated, scramble and GID/CTLH knockout (*MAEA*^-/-^) BMDM^h^s with the wild type *Mtb* Edman strain at MOI 0.3 in similar conditions as outlined in Fig. 1. On day 4, infected macrophages were treated with INH (0.4 μg/ml) or Pa (1.3 μg/ml) for 4 days. CFUs were plated from untreated infected macrophages on day 0 (3 hours post infection), day 4 (on the day of drug addition) and after 4 days of drug treatment on day 8. Consistent with our luciferase data (Fig. 2, Fig. 4), both LPS/IFNγ activated and *MAEA*^-/-^ BMDM^h^s significantly controlled *Mtb* growth as evidenced by reduced number of CFUs when compared to resting or scramble macrophages respectively after 4 days of infection (Fig. 6A, 6B). CFU numbers were comparable and not significantly different on day 0, 3-hours post-infection (Fig. 6A, 6B). After 4 days of drug treatment, we observed that *Mtb* exhibited reduced susceptibility to INH in both LPS/IFNγ activated and *MAEA*^-/-^ BMDM^h^s when compared to controls (Fig. 6A, 6B). Even though Pa had comparable efficacy in scramble and *MAEA*^-/-^ BMDM^h^s (Fig. 6B), the drug displayed increased potency against *Mtb* in LPS/IFNγ activated BMDM^h^s as indicated by lower CFU numbers when compared to resting macrophages (Fig. 6A). Together, these data correlated strongly with our luciferase measurements (Fig. 2A, Fig. 4A, Fig. 3C, Fig. 5B) adding further validity to the robustness of the assay.

**Fig. 6:**
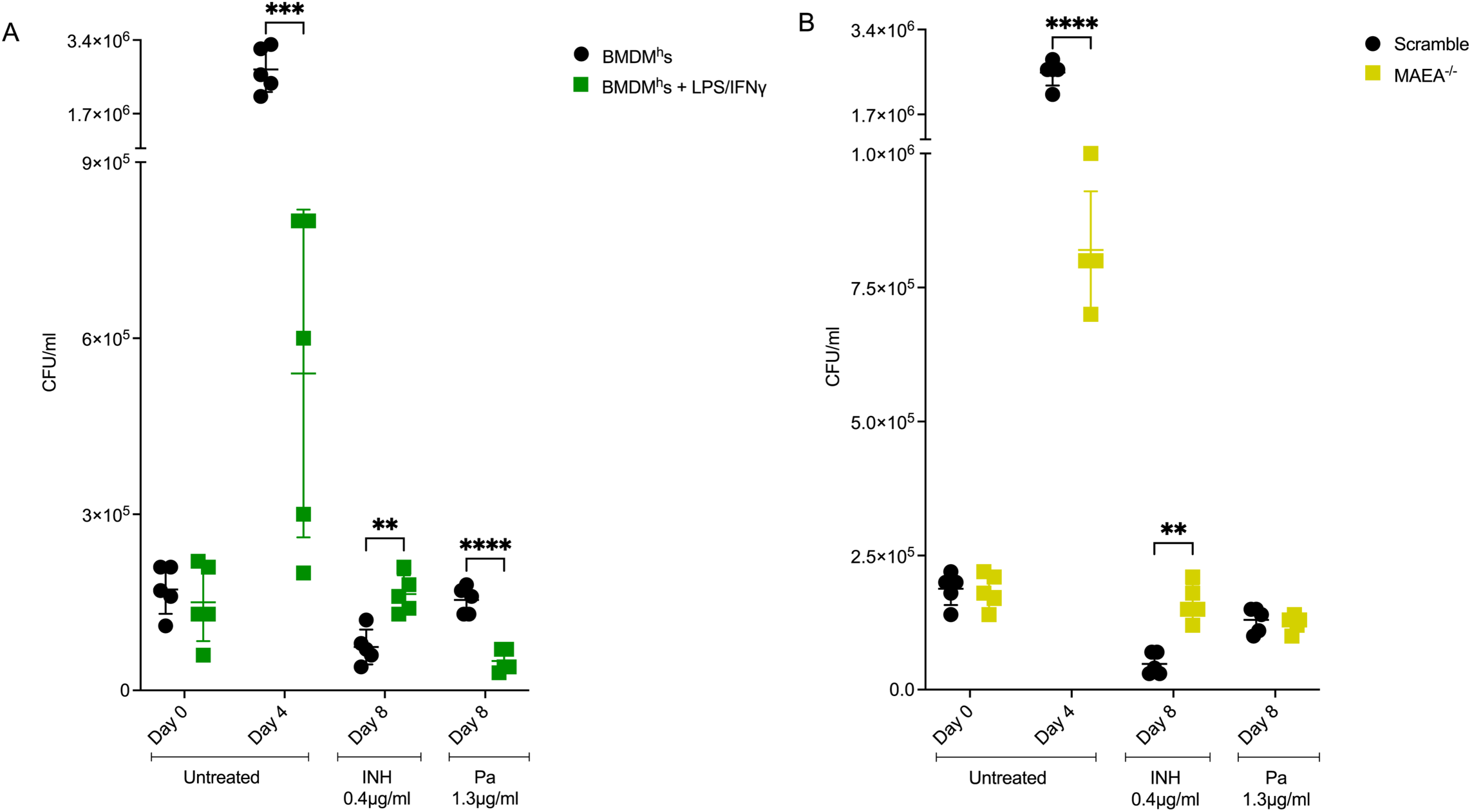
M*t*b CFU measurements in non-treated and drug treated controls, LPS/IFNγ activated and *MAEA*^-/-^ BMDM^h^s. Resting, LPS/IFNγ activated, scramble and MAEA^-/-^ macrophages were infected with *Mtb* Erdman at MOI 0.3. Cells were left untreated or treated from day 4 with INH 0.4 μg/m or Pa 1.3 μg/ml. A. Plated CFUs in resting and LPS/IFNγ activated BMDM^h^s for the indicated days and treatment conditions. B. Plated CFUs in scramble and *MAEA*^-/-^ BMDM^h^s for the indicated days and treatment conditions. n=5 biological replicates; ** P < 0.01, ***P < 0.001, **** P < 0.0001. two-way ANOVA alongside Dunnett’s multiple comparison test. Data are presented as mean values ± SD.

## Conclusion

In summary, we present a new 96 well plate, macrophage infection assay that enables one to determine the impact of host cell physiology on the efficacy of anti-mycobacterial drugs. *In vivo* and intracellular environments present a significant challenge to the development of new therapeutics with regards to drug pharmacokinetics, stability and penetrance. These problems are amplified in TB therapeutics where treatment must always involve the use of multiple drug combination therapies. In the development of new tools to address these challenges, we feel that one area that has lagged behind is the development of *in vitro* assays that reliably captures the biology of differential drug susceptibility of *Mtb in vivo* and within the host cell. The assay we have developed relies on induction of host stresses on intracellular bacteria either by cytokine activation or CRISPR-Cas9 knockout of host genes as surrogates for the immune environments the bacteria likely encounters *in vivo*. Following drug exposure in the different host cell environments in the assay platform, the relative magnitude and fitness of the surviving bacterial population is assessed by outgrowth after host cell lysis. Measurement of luciferase signal facilitates resolution of small but significant differences in differential drug susceptibility. We believe that the ability to profile existing and emergent anti-TB drugs for their potency against *Mtb* in different physiological states of *in vivo* relevance, as our assay demonstrates, is of particular significance in designing drug regimens with improved coverage of the heterogeneous *Mtb* subpopulations *in vivo* [18].

## Materials and methods

### Cell culture conditions

Wild-type (C57B6/J mice) and Cas9^+^ (Jackson stock 026179) Hoxb8 conditionally immortalized myeloid progenitor cell lines derived from murine bone marrow were generated according to the protocol previously described [19]. Hoxb8 myeloid precursors were cultured in RPMI (Corning^®^) media supplemented with 10% FBS (fetal bovine serum), 2 mM L-Glutamine, 1 mM Sodium pyruvate, 20 ng/ml murine GM-CSF (PeproTech), 10mM HEPES, 0.5 μM β-estradiol and 1 % penicillin/streptomycin. BMDM^h^s were differentiated from the myeloid progenitors by washing twice with PBS to remove any remnants of β-estradiol and seeding in BMDM differentiation media; DMEM (Corning^®^) supplemented with 10% FBS, 15% L929 cell conditioned media, 2 mM glutamine, 10 mM HEPES, 1 mM sodium pyruvate and 1 % penicillin/streptomycin for 6-7 days. To perform the intracellular drug assay under immune activation, BMDM^h^s were activated with media containing 20 ng/mL of recombinant mouse IFNγ (Peprotech) and 100 ng/ml LPS 24 hours prior to infection. 20 ng/mL IFNγ was maintained in macrophage media throughout the experiment after infection. GID/CTLH knockout BMDM^h^s were generated as described previously [21].

### Bacterial strain and antibiotic stocks

The *Mtb* Erdman *strain* (ATCC 35801) was used in CFU experiments. Luciferase expressing *Mtb* Erdman*-*Lux strain was generated by the transformation of the pMV306G13+Lux plasmid [55]. Bacterial strains were grown to logarithmic phase at 37°C in Middlebrook 7H9 broth supplemented with 10% OADC (oleic acid-albumin-dextrose-catalase), (Becton Dickinson and Company), 0.2% glycerol, 0.05% tyloxapol (Sigma-Aldrich) and 25 μg/ml Kanamycin (*Mtb* Erdman*-* Lux strain). All antimicrobial agents used in the study (INH, RIF, EMB, PZA, BDQ, LZD, Pa and MOX) were purchased from MedChemExpress. Stock solutions of INH, EMB and PZA were prepared in water. Stock solutions of RIF, BDQ, LZD, Pa and MOX were prepared in DMSO.

### Macrophage infections

BMDM^h^s were seeded in 96-well (1 x 10^5^ cells/well), white optical-bottom plates (Thermo Scientific) and the infection with *Mtb* was performed as previously described [21, 56]. Briefly, BMDM^h^s were seeded overnight at 37°C in 5% CO_2_ before infection. Log phase bacteria were pelleted, resuspended in basal uptake buffer (2.25 g glucose, 2.5 g bovine serum albumin, 0.5 ml gelatin, 50 mg CaCl_2_, 50 mg MgCl_2_, 500 ml 1x PBS) and syringed for around 15-20 times using a BD 3 ml luer-lock syringe with 25-gauge needle. The bacterial suspension was resuspended in macrophage media without antibiotic to achieve a MOI of 0.3 or 0.4 and then added to the macrophage monolayers. Luminescence was measured on an Envision plate reader (PerkinElmer) and the bacterial growth kinetics were monitored throughout the assay over 12 days.

### CFU enumeration of *Mtb* in macrophages

Confluent macrophage monolayers were infected with *Mtb* at MOI 0.3. 3-hours post infection, extracellular bacteria was removed by washing with fresh macrophage media at least 3 times. At various time points, macrophages were lysed with 0.01% SDS in H_2_0 for 15 minutes and diluted in 0.05% Tween-80 in 1x PBS. Diluted lysates were plated on 7H10 OADC agar plates and incubated at 37^0^C for at least 3 weeks.

### Data analysis

Growth kinetics graphs were constructed based on measured RLUs with subtracted background from uninfected cells. Plots and statistical analyses were performed in Graphpad prism (v 10.4.0).

## Acknowledgements

We would like to thank Dr. Chris Sassetti at UMass Chan Medical School who provided the Hoxb8 cell lines used in the study. This work was supported by grants from the National Institutes of Health (AI155319, AI162598, and OD032135), Bill and Melinda Gates Foundation and the Mueller Health Foundation to D.G.R and a postdoctoral seed grant to N.V.S. from the Cornell Center for Antimicrobial Resistance Research and Education.

